# Conjugative plasmid transfer is limited by prophages but can be overcome by high conjugation rates

**DOI:** 10.1101/2021.03.29.437513

**Authors:** Claudia Igler, Lukas Schwyter, Daniel Gehrig, Carolin Charlotte Wendling

## Abstract

Antibiotic resistance spread via plasmids is a serious threat to successfully fight infections and makes understanding plasmid transfer in nature crucial to prevent the rise of antibiotic resistance. Studies addressing the dynamics of plasmid conjugation have yet neglected one omnipresent factor: prophages (viruses integrated into bacterial genomes), whose activation can kill host and surrounding bacterial cells. To investigate the impact of prophages on conjugation, we combined experiments and mathematical modelling. Using *E. coli*, prophage *λ* and the multidrug-resistant plasmid RP4 we find that prophages can substantially limit the spread of conjugative plasmids. This inhibitory effect was strongly dependent on environmental conditions and bacterial genetic background. Our empirically parameterized model reproduced experimental dynamics of cells acquiring either the prophage or the plasmid well but could only reproduce the number of cells acquiring both elements by assuming complex interactions between conjugative plasmids and prophages in sequential infections. Varying phage and plasmid infection parameters over empirically realistic ranges revealed that plasmids can overcome the negative impact of prophages through high conjugation rates. Overall, the presence of prophages introduces an additional death rate for plasmid carriers, the magnitude of which is determined in non-trivial ways by the environment, the phage and the plasmid.

## Introduction

Mobile genetic elements (MGEs), such as plasmids and bacteriophages move horizontally between bacterial cells and thereby represent a key source of microbial genetic diversity. Conjugative plasmids (i.e., plasmids that mediate their own transmission via conjugation) frequently carry accessory genes beneficial for their hosts, which can, for example, confer resistance to heavy metals or antibiotics [1, 2]. Predicting plasmid dynamics and the horizontal transfer of their accessory genes is therefore not only important for understanding bacterial ecology and evolution in general, but also for anticipating the rise of antibiotic resistance.

Accordingly, conjugative plasmid transfer has been a topic of intense research, revealing a complex, tightly regulated process (reviewed in [3]). Most of these mechanistic studies were necessarily conducted in well-engineered environments, which allow the investigation of plasmid conjugation in isolation. Yet, wherever plasmids and their host bacteria exist, they are generally surrounded by a diverse repertoire of other MGEs, particularly bacteriophages [4]. Earlier work on plasmids and bacteriophages frequently showed strong interactions between them, mostly resulting in interference, either one way (plasmid inhibition of phage reproduction [5, 6]), or bi-directional, leading to diminished phage and plasmid reproductive success [7, 8]. The former was for instance due to specific anti-phage systems (restriction-modification systems [9]), whereas the latter resulted from sharing of host resources [8]. The high number of plasmids that contain general mechanisms for inhibiting phage infections (i.e., effective against many different phage species [5, 6]) indicates a benefit for plasmids to protect their hosts against phage infections.

Why would plasmids need to defend themselves and their host cells against phages? For one, bacteriophages are a major cause of bacterial mortality through cell lysis [10, 11]. These strong reductions in bacterial densities can limit plasmid persistence and spread, and even lead to plasmid extinction [12]. In other cases, plasmid-phage interactions were found to be more specific: for instance, filamentous phages, which do not lyse their host cells but usually establish a chronic infection therein, use conjugative pili as receptors, which can inhibit plasmid transfer [13]. The impact of cell death via obligately lytic phages (i.e., infection always results in cell lysis) [12] and specific mechanisms of interference on plasmid conjugation seem straight-forward. What remains unclear is how temperate phages, which – similarly to conjugative plasmids – can be transferred both vertically and horizontally, affect conjugative plasmid transfer. Upon infection of a bacterial host cell, temperate phages can choose between a lytic or a lysogenic life cycle. At a low ratio of phages to bacterial cells they mainly go through lytic infections, whereas higher ratios increase the likelihood that temperate phages choose the lysogenic cycle [14–16]. By integrating their own genetic material into the bacterial genome, the phage becomes a prophage and the infected host bacterium a lysogen. Prophages can switch to the lytic cycle to replicate, assemble progeny and release phage virions via host cell lysis [17], either spontaneously (at low rates) or in response to stressful conditions (at high rates, [16, 18]). As lysogens are usually immune against superinfection by the same phage [19, 20], released phage particles can only productively infect and potentially kill competing non-lysogens, which are susceptible to these phages.

Due to their complex lifestyles, we predict that temperate phages are likely to influence plasmid transfer dynamics in nature in ways less obvious as for lytic phages [12]. Similar to lytic phages, free or induced temperate phages can kill plasmid-carrying and recipient cells [21], which will negatively influence plasmid spread and persistence [12]. On the other hand, resident prophages will protect plasmids carried in the same host cell from lytic infections due to superinfection immunity [19, 20]. By focusing on these indirect effects of prophages on plasmid transfer dynamics – instead of specific interference mechanisms [9] – we aim to unravel general interaction patterns of temperate phages and plasmids. Particularly, we are interested in how such phage-plasmid interactions are shaped by their infection characteristics, such as transfer rates, or by environmental conditions.

To determine how the presence of an active prophage influences plasmid transfer dynamics, we performed conjugation experiments in two environments that differ in their phage adsorption rates. For the experiments we used *E. coli*, its naturally associated prophage *λ* and the conjugative multidrug-resistant plasmid RP4. Additionally, we used a mathematical model to identify crucial phage and plasmid infection parameters, which we then varied over a wide range that captures empirically reported values and explores the generality of our results. The model could reproduce phage and plasmid acquisition under all experimental conditions well by assuming differential infection of empty and MGE-carrying hosts, indicating unknown prophageplasmid interactions that deserve further exploration. Overall, we find that prophages can limit plasmid spread substantially by effectively introducing an additional (density-dependent) death rate, which can however be overcome by higher plasmid conjugation rates.

## Methods

### Experimental work

#### Strains,media and culture conditions

All experiments were carried out in liquid LB (Lysogeny Broth, Sigma-Aldrich) (‘LB’) or liquid LB supplemented with 10% SM Buffer (9.9 mM NaCl, 0.8 mM MgSO4 × 7 H2O, and 5 mM Tris-HCl) (‘SM’). Antibiotics were used at the following concentrations, unless otherwise noted: Ampicillin (amp): 100 µg mL^-1^; kanamycin (kan): 50 µg mL^-1^, tetracycline (tet) and chloramphenicol (cm): 25 µg mL^-1^. Mitomycin C was used at a concentration of 0.5 µg mL^-1^. All bacterial strains used in the present study were designed from wildtype *E. coli* K-12 MG1655 (WT; Table 1). We used three different donor strains: (1) a plasmid donor (‘independent plasmid donor’), which carried the RP4 plasmid conferring resistance to three different antibiotics: ampicillin (amp^R^), kanamycin (kan^R^), and tetracyclin (tet^R^), (2) a phage donor (‘independent phage donor’), i.e., a lysogen carrying the prophage *λ* which encoded an ampicillin resistance gene, and (3) a plasmid phage donor (‘common donor’), which carried both, the RP4 plasmid and the *λ* prophage. Lysogen construction was done as described in [21]. In addition, the independent phage donor and the common donor, were both labelled with a yellow-super-fluorescent protein (SYFP) marker [22]. The recipient was labelled with a dTomato fluorescent protein and carried a chloramphenicol (cm^R^) resistance gene (kindly provided by Lei Sun and Erik Gullberg). Strains were grown in the LB or SM environment at 37^*°*^C with constant shaking at 180 rpm.

**Table 1:** Strains and phage used in the present study.

| Strain/Phage | Genetic background |
| --- | --- |
| Wildtype | *E. coli* K-12 MG1655 |
| Independent plasmid donor | RP4 plasmid (amp<sup>R</sup>, kan<sup>R</sup>, tet<sup>R</sup>) |
| Independent phage donor | SYFP, λ+ Δ(bor)::amp<sup>R</sup> |
| Common donor | SYFP, λ+ Δ(bor)::amp<sup>R</sup>, RP4 plasmid (amp<sup>R</sup>, kan<sup>R</sup>, tet<sup>R</sup>) |
| Recipient | dTomato (cm<sup>R</sup>) |
| λ+ | Δ(bor)::amp<sup>R</sup> |

#### Bacterial and phage densities

Total bacterial counts were determined by plating out 25 µl of a dilution series ranging from 10^−1^ to 10^−6^ on LB-agar plates (Sigma-Aldrich). To identify recipient strains (carrying the dTomato marker) we took pictures of each plate using a ChemiDoc XRS+ and subsequently counted the colonies using ImageJ with the PlugIn CellCounter for manual cell count. To determine phage donors (carrying the SYFP marker) we used a blue light. Further discriminations between all possible strains and prophage-plasmid combinations were achieved by replica plating (details are described below for each experiment separately). All plates were incubated overnight at 37^*°*^C and the total amount of colonies was counted the following day. The amount of free viral particles (PFU/mL) per culture was determined using a standard spot assay, where 2 µl of a dilution series of phages ranging from 10^−1^ to 10^−8^ diluted in SM-Buffer was spotted onto a bacterial lawn containing 100 µl indicator cells of the WT grown in 3 mL of 0.7% soft-agar. To obtain indicator cells we centrifuged 15 mL of an exponentially growing WT culture for 10 min at 5000 rpm, resuspended the pellet in 2 mL 10mM MgSO4 and incubated the mixture for another hour at 37^*°*^C with constant shaking.

#### Experimental design

We used two different single-donor experiments to determine conjugation-and lysogenization-(i.e., lysogen formation, which subsumes various processes including phage induction, adsorption and life cycle decision making, i.e., lysogeny vs. lysis) rates in LB and SM respectively. To better parameterize our model, we performed two additional MGE-transfer experiments, which estimated whether the presence of one MGE in the recipient influences the transfer of the other MGE. To do so we co-cultured either the independent phage donor with the recipient that carried the plasmid, or vice versa. Note, that both MGEs (the one from the recipient and the one from the donor) are capable of horizontal transfer in these experiments.

These transfer control experiments were then followed by an experiment in which we cocultured the recipient with both independent-donor strains (referred to as independent-donor experiment), to investigate how the presence of active prophages influences plasmid dynamics and how this depends on the environment (LB or SM). To favour plasmid spread, we performed a third experiment in which we co-cultured the recipient with the common donor that harboured the plasmid and the prophage simultaneously (referred to as common-donor experiment). In this experiment, we additionally tested the model prediction that different starting cell-densities do not influence transfer dynamics in the presence of LB and SM. Each experiment was done with three to four (single-donor experiments) or six (common-and independent-donor experiments) independent biological replicates. For each replicate, single colonies of each strain were inoculated in 6 mL of LB and grown overnight. The following day we determined the optical density of each overnight culture at 600 nm using an automated plate-reader (Tecan infinite 200) and adjusted the OD of each culture to the lowest measured OD, to obtain similar starting concentrations of each subpopulation per culture.

Note, that while we do not add free phages in any of the experiments, there will likely be some free phages carried over from the diluted overnight lysogen cultures, stemming from spontaneous induction. Using our model (considering spontaneous induction and loss due to phage decay or adsorption to lysogenic cells), we calculated this number to be between 10^4^ and 10^6^ phages (two orders of magnitude below the lysogen starting density), depending on the magnitude of the dilution.

##### Single-donor experiments

To determine the conjugation rate of RP4 or the lysogenization rate of phage *λ*, we diluted the OD-adjusted overnight cultures of donors (either the phage or the plasmid donor) and recipients 1:30 in each of the two environments and subsequently added 300 µl of each dilution to 5400 µl LB or LB+SM, respectively. This resulted in a starting concentration of approximately 2×10^6^ CFU/mL of each strain. We then determined PFU/mL and CFU/mL from seven time-points, i.e., T0 immediately after mixing, T1-T7 corresponding to 1, 2, 3, 5, and 7 hpm (hours post mixing) and T24 at 24 hpm. CFU/mL was determined by selective plating on LB to estimate the total population density and LB supplemented with cm and kan to select for transconjugants (recipient plasmid carriers) or on LB supplemented with cm and amp to select for new lysogens (recipients carrying the prophage).

##### MGE-transfer control experiments

We used a similar approach as for the single-donor experiments to determine conjugation and lysogenization in the presence of the respective other MGE with the following alterations: To better compare the dynamics to the independent-donor experiment we used the same starting cell densities as therein, i.e., 3×10^7^ CFU/mL. CFU/mL were determined as describe above by selective plating at T0, T1, T3, T7 and T24. Phage-acquisition by plasmid carriers was determined by a mitomycin C assay on a random set of 24 amp-resistant recipient or donor clones followed by a spot assay on the phage-susceptible wildzztype. Recipients and donors were discriminated based on their flurorescence marker (SYFP marker: phage or plasmid donor, dTomato: recipient).

##### Independent-donor experiments

To determine how prophages influence plasmid dynamics we co-cultured both donor strains and the recipient strain at equal starting concentrations of 3×10^7^ CFU/mL of each strain type. OD-adjustment and dilutions were done as described above for the single-donor experiments. Plating at selected time points (T0 – T24, as described for the single-donor experiments) was done on (1) LB-agar to obtain total community densities, (2) LB- agar + kan to select for all plasmid-carrying strains, (3) LB-agar + cm to select for all recipients. Subsequently LB-agar + cm plates were stamped on three different selective agar plates: (3.1) LB as a reference for total cell count, (3.2) LB-agar + amp to select for recipient strains that have acquired the phage, the plasmid or both, (3.3) LB-agar + kan to select for recipient strains that have acquired the plasmid. Colonies from plates (1) and (2) were further discriminated based on the fluorescent marker. In addition, we screened a subset of amp resistant recipients (n=24) for phage presence using a mitomycin C induction followed by a spot-assay on the WT to estimate the proportion of lysogens and transconjugants, which are both resistant to ampicillin.

To determine whether resistance against the ancestral phage had emerged we each picked 40 colonies from recipients and plasmid donors from every replicate after 24 hours. From each colony we generated indicator cells to be used in a standard spot assay in which we spotted 2 µl of the ancestral phage on a 3 mL lawn containing 100 µl indicator cells. No plaque formation indicated resistance evolution, but no heritable resistance was found at the end of the experiment.

##### Common-donor experiments

We favoured conjugation using a common donor, carrying the phage and the plasmid, and tested that final transformant cell numbers were relatively independent of starting densities, by performing a separate experiment using four different environments: LB/high, LB/low, SM/high, SM/low, where high and low correspond to the total population density at the onset of the experiment (high: ∼6.7×10^8^ CFU/mL; low: ∼4.6×10^6^ CFU/mL). The common plasmid phage donor should give the plasmids an additional advantage through prophage-mediated immunity to further phage infections, thereby avoiding the heavy (initial) lysis of plasmid donors observed in the independent-donor experiment. OD-adjustments and dilutions were done as described above and the total populations were plated on selective plates at T0, T2, T5, and T24 hpm. We discriminated the different colony types by selective and replica plating on LB-agar plates carrying different antibiotics and based on their fluorescent marker as described for the independent-donor experiment. We expect colonies growing on plates containing cm, kan and amp to be recipients that have acquired the plasmid and potentially the phage. To further discriminate between transconjugants and double-carriers we screened a subset of those colonies (n=24) for phage presence using mitomycin C induction followed by a spot-assays on the WT.

##### Estimating bacterial growth rates

To better parameterize our model, we measured bacterial growth rate of our donor and recipient strains with and without the phage, the plasmid or both elements. Briefly, overnight cultures were diluted 1:100 into fresh LB and grown for 24 hours in a plate-reader. Optical density was measured every 10 minutes at 600 nm. Growth rates were determined by calculating doubling times from exponential growth data. As especially lysogen cultures not always showed strictly exponential growth, doubling times were calculated five times for slightly different parts of the exponential growth curve (varying start and end points by 15 to 45 min) and the mean of those calculations was taken as growth rate for the model.

##### Estimating phage latent period

Phage latent period was estimated from the rise of one-step growth curves performed in triplicates. Overnight cultures of the WT were inoculated 1:100 into fresh LB and grown for another four hours to mid-exponential phase. Afterwards 1 mL of bacteria were mixed with approximately 10^6^ pfu/mL (leading to an MOI≤1) and phages were allowed to adsorb for 20 min at 37° C without shaking. The mixture was then diluted 1:100 into fresh LB and incubated at 37°C shaking. The increase in phage density was repeatedly determined from the mixture at 10-minute intervals using a standard spot assay on lawns of the WT.

#### Statistical analysis

##### Detection limit

To prevent overgrowing of plates we had to adjust the dilutions that we chose to plate out to be able to count CFUs for the total population density (which was several orders of magnitude higher than of individual fluorescent subpopulations). By doing so, we introduced a detection limit for several subpopulations, which, depending on the total population size, varied for the different time-points. In order to account for this detection limit, we calculated the mean between the lowest dilution (that we plated and that still had countable colonies for each timepoint and plate-type) and zero, log-transformed this value and assigned it to the respective subpopulations.

##### Differences in subpopulation densities

To determine differences between subpopulations across experiments, treatments or time points, we used linear models followed by pairwise comparisons using Tukey’s HSD (R package emmeans) to account for multiple testing. While we generally distinguish between cells carrying one or both MGEs, statistical tests were performed with ‘total’ transconjugants (transconjugants plus double-transformants) or ‘total’ lysogens (lysogens plus double-transformants), in order to make them comparable with single-donor experiments.

### Mathematical model of phage and plasmid transfer dynamics

We used a mathematical model to describe the dynamics of individual bacterial subpopulations (bacterial cells carrying one or both or neither MGE) as well as the phage population, making the following assumptions: Plasmid transfer and phage infections can be described via mass action kinetics, as experimental cultures were grown in liquid medium with shaking, providing a well-mixed environment. As further all entities are generally found at high numbers, we used a deterministic model. Phage and plasmid infection dynamics lead to significant, but ‘fixed’ time delays (due to, either the time spent within the cell for multiplication, or the time necessary for DNA transfer), hence we used delay-differential equations. The model was implemented in Matlab (The MathWorks, Natick, MA, USA)), using the dde23 solver, as were the algorithms for fitting parameters (simulated annealing), model comparisons (RSS and AIC) and parameter sensitivity analysis (see below). The simulation time for population dynamics was 24h, using starting densities and parameters fitted to empirical data or taken from literature (see ‘Parameter values and estimation’; Table S1-S5).

#### Independent-donor model

(Figure 1E, equations (10)-(24), Table S1&S2)

**Figure 1:**
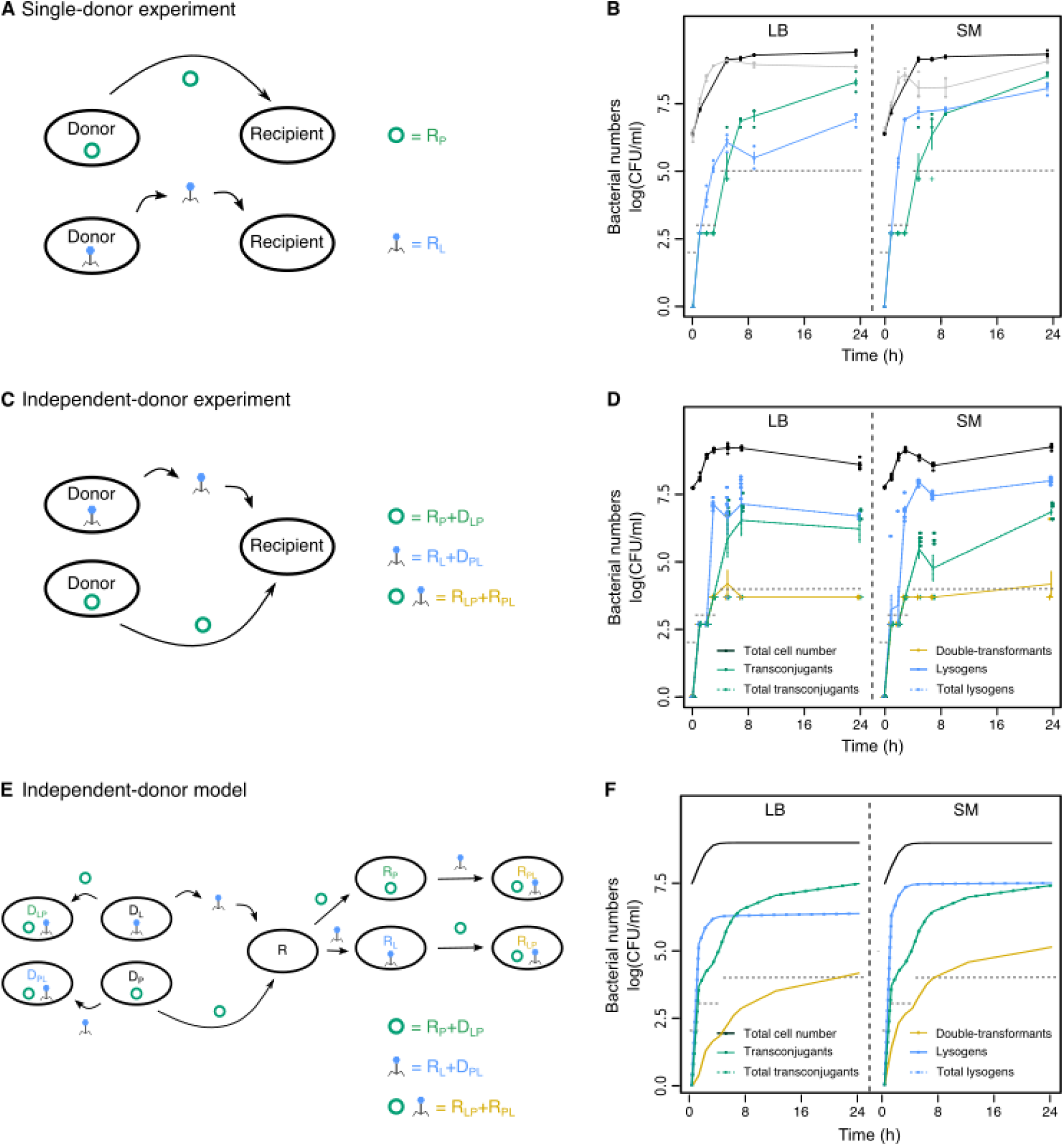
Phage and plasmid transfer with independent donors. (A) Single-donor experiments with independent plasmid and phage transfer. (B) Total amount of bacteria in conjugation (black) or lysogenization (grey) experiments, newly formed transconjugants (*R*_*P*_ ; green) and newly formed lysogens (*R*_*L*_; blue) measured as colony forming units log(CFU/mL) over time in LB (left panel) or SM (right panel) in single-donor experiments (shown are single data points as well as mean ± s.e., n=4). Grey dashed lines indicate the empirical detection limit at a given time point (Methods). (C) Independent-donor experiments with one donor carrying the plasmid (*D*_*P*_) and one carrying the phage (*D*_*L*_). Transconjugants (green) are counted as recipient and lysogen donors, which acquired the plasmid (*R*_*P*_ + *D*_*LP*_), and lysogens (blue) as recipients and plasmid donors, which acquired the phage (*R*_*L*_ + *D*_*P L*_). Double-transformants (yellow) are recipients, that acquired the plasmid and the phage (*R*_*LP*_ + *R*_*P L*_). (D) Total cell number (black) as well as newly formed lysogens (blue), transconjugants (green), and doubletransformants (yellow) measured as colony forming units log(CFU/mL) over time in LB or SM in independent-donor experiments (shown are single data points as well as mean ± s.e., n=6). For comparison with (B) total transconjugants (*R*_*P*_ + *D*_*LP*_ + *R*_*P L*_ + *R*_*LP*_) and total newly formed lysogens (*R*_*L*_ + *D*_*P L*_ + *R*_*P L*_ + *R*_*LP*_) are shown in dashed green or dashed blue lines, but are hidden by the respective solid lines. (E) Model reactions and (F) 24h-simulations of phage and plasmid transfer for the independent-donor experiment (colours and line types as described in C&D; see Methods for model details).

##### Growth

Bacterial cells grow at a maximal net growth rate *r* (summarizing growth and death) until they reach carrying capacity *K*:

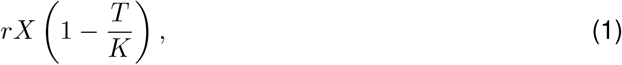

where *X* here indicates any of the bacterial subpopulations described below and *T* the total number of bacterial cells (Equation 23) summed over all subpopulations (i.e., wildtype, phage-carrying and plasmid-carrying subpopulations, which are defined below). Cells carrying no MGE (recipient *R*), the plasmid (*X*_*P*_), the prophage (*X*_*L*_) or both (*X*_*LP*_, *X*_*P L*_) grow at *r, r*_*p*_, *r*_*l*_ and *r*_*lp*_ respectively, using empirically determined growth rates (S5; see ‘Estimating bacterial growth rates’). Note, that we here differentiate subpopulations with regard to the order of MGE acquisition (as we are interested in the transfer dynamics), but the growth rate of cells carrying both MGEs, *r*_*lp*_, is independent of acquisition order.

##### Conjugation

Plasmids are transferred from any of the plasmid-carrying populations (which are summed into *P*) to recipient cells (*R*), phage donors (*D*_*L*_) or recipient lysogens (*R*_*L*_) with conjugation rate *p*. (Initially, plasmids are only present in our system in the plasmid donor subpopulation, *D*_*P*_ .) After *τ*_*p*_ time units needed for transfer, conjugation results in recipient plasmid (*R*_*P*_), phage donor plasmid (*D*_*LP*_) and recipient lysogen plasmid (*R*_*LP*_) cells, respectively:

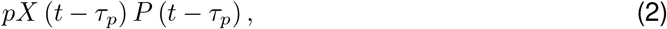

with *X* ∈ {*R, D*_*L*_, *R*_*L*_} being one of the non-plasmid populations, and *P* = *D*_*P*_ + *R*_*P*_ + *D*_*LP*_ + *D*_*P L*_ + *R*_*P L*_ + *R*_*LP*_ the entirety of all plasmid-carrying cells.

Our empirical data suggested that conjugation slows down when cells approach stationary phase (see also [23, 24]), but is not completely inhibited (see increase of transconjugants from T7 to T24 in Figure 1B), hence we used a logistic function for conjugation, which is scaled by *k*_*p*_:

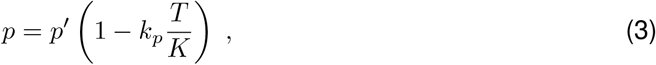

where *p*^*′*^ is the maximal conjugation rate.

Fitting our parameters to empirical data (see ‘Parameter values and estimation’) indicated that plasmid conjugation occurred at different rates, *p*_*l*_, to lysogens, which was 10-fold lower than to wildtype recipients in the independent-donor experiments.

Including segregation loss of plasmids seemed to be negligible in this system and did not improve our model fit with empirical data.

##### Phage infection

Free phage virions (*V*) can infect recipient cells (*R*), plasmid donors (*D*_*P*_) and recipient plasmid carriers (*R*_*P*_) at rate *α* (adsorption rate), which results either in (1) lysogeny, i.e., integration of the phage genome into the host genome, producing lysogen cells (*R*_*L*_, *D*_*P L*_ and *R*_*P L*_ respectively), or (2) lysis, i.e., productive infection, which leads to host cell lysis and release of *β* new virions. In non-lysogen cells (lysogens are protected from phage infection by superinfection immunity), infecting phages take the lytic pathway at rate (1 − *l*) and cells are lysed after a latent period of *τ*_*l*_ time units:

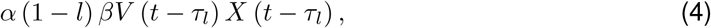

for *X* ∈ {*R, D*_*P*_, *R*_*P*_}.

Similarly, phages infecting non-lysogen cells can make a decision for lysogeny (after *τ*_*d*_ time units) with probability *l*:

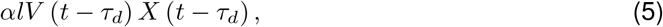

for *X* ∈ {*R, D*_*P*_, *R*_*P*_}.

As has been shown before [14], the lysogenization probability is dependent on the number of phages infecting a bacterial host cell and can, in our well-mixed system, be approximated by a linear function of the number of free phages:

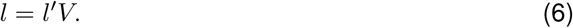

Integrated phages can spontaneously switch to the lytic infection mode again (induction), which we assume to occur at a low constant rate *i* ([23, 24]). This leads to host cell lysis and the production of *β* new virions after *τ*_*l*_ time units:

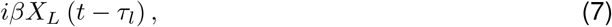

for *X* ∈ {*D, R, D*_*P*_, *R*_*P*_}.

The process of phage infection is usually modelled in a simplified manner via a single, constant parameter, the adsorption rate *α* [25]. This parameter is however quite complex as it summarizes several biological processes necessary for successful phage infection, like extracellular diffusion, attachment to a host cell and phage genome injection. For many phages the adsorption rate *α* is dependent on the physiological state of the host bacteria and saturates with *K* [26, 27], which, according to our empirical data, is also true for phage *λ* (Figure 1B). Using a logistic function for the growth-dependence of *α*, we found that the best fit to our data came from a steep response to the approach of stationary phase, which we included via a power-law exponent (*k*_2_), but – similarly to the growth rate dependence of the conjugation rate – not a complete inhibition at carrying capacity (as scaled by *k*_*a*_):

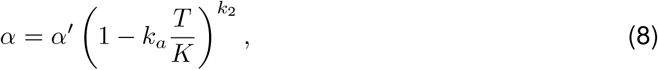

where *α*^*′*^ is the maximal adsorption rate.

Free phages ‘die’ either through decay at rate *γ*_*V*_, or they are ‘lost’ by infection of lysogen cells, which cannot be productively infected:

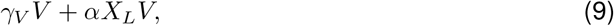

for *X* ∈ {*D, R, D*_*P*_, *R*_*P*_}.

##### Initial conditions

Model simulations were started at bacterial densities of recipients and donor strains as used to inoculate the experiments. Further, we started with a free phage density that was two orders of magnitude lower than the lysogen starting density to account for the carry- over effect of free phages from overnight lysogen cultures. We calculated this number based on spontaneous phage induction, phage loss due to adsorption to lysogen cells and phage decay, from an overnight culture of phage donors and subsequent dilution of cells (and free phages) to a given initial bacterial starting density used in the respective experiments.

The processes described above led to the model equations shown below. (For better readability, we do not differentiate between growth rates of lysogens (*r*_*l*_), plasmid carriers (*r*_*p*_), doubletransformants (*r*_*lp*_), depending on the bacterial donor or recipient background in the equations below, but the values inferred through fitting to empirical data are given in Table S1&S2.) Here, we model plasmid and phage transfer as a sequential process, but including the possibility of a simultaneous transfer to recipients did not change our results notably due to the low probability of these events.

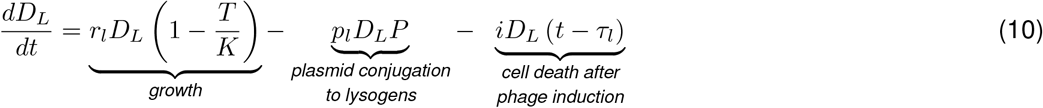

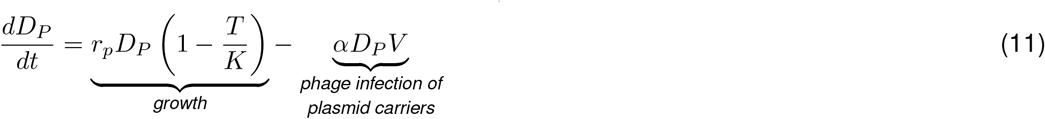

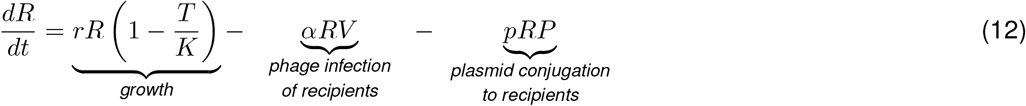

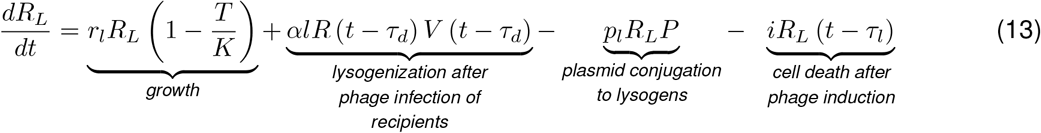

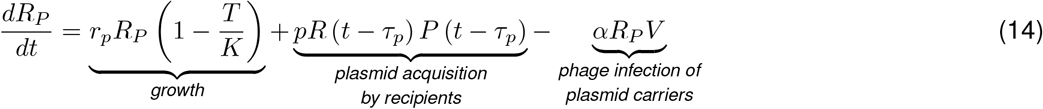

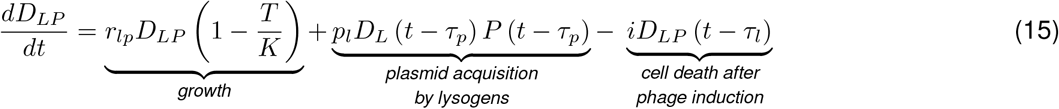

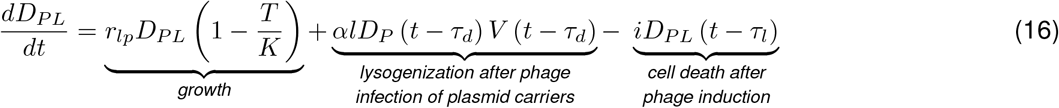

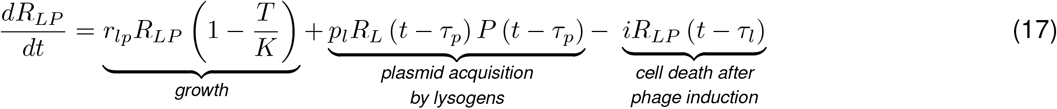

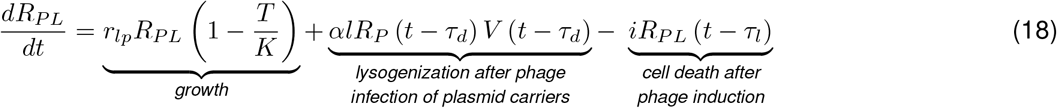

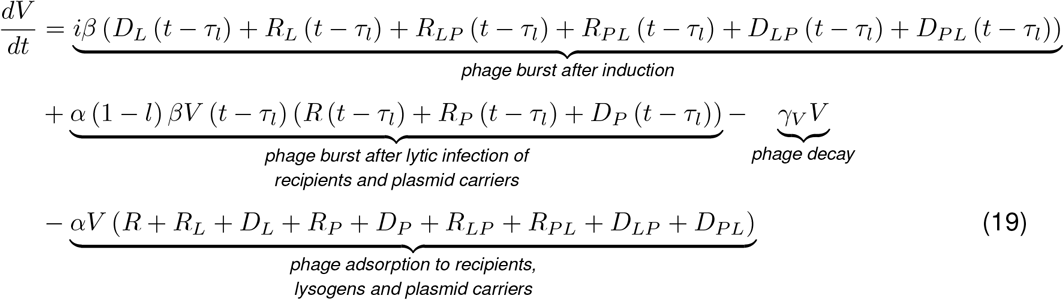

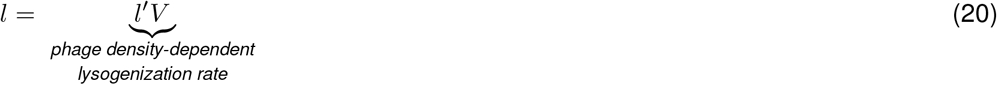

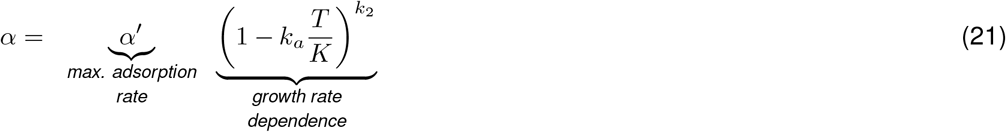

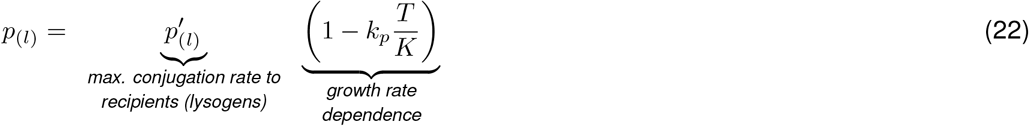

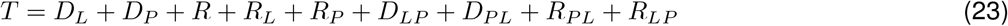

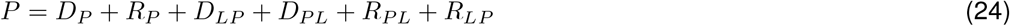

##### Parameter values and estimation

#### Directly estimated parameters

Some of the model parameters could be determined directly from empirical data: Bacterial growth rates of donor and recipient cells carrying no, one or both MGEs (*r, r*_*p*_, *r*_*l*_, *r*_*lp*_) were determined via linear regression as the slopes of growth measurements for each individual population, giving effective growth rates, i.e., including cell growth and death (Table S5). Carrying capacity was determined from stationary phase CFU counts of all bacterial cells in conjugation experiments over time (Figure 1B).

Latent period, *τ*_*l*_, was determined from phage infection experiments (one-step growth curves) as the time until phage numbers started to increase (minimal latent period), which was very similar to reported values [28].

#### Fitted parameters

Most infection parameters were fitted to empirical data by using ‘simulated annealing’ [29]. Briefly, the neighborhood of a given parameter set is explored for a better simulation fit, calculated as the difference (Residual Sum of Squares) between the true observation points and the simulated ones. A parameter set is then accepted dependent on its goodness of fit, but in a probabilistic manner. In this way the algorithm avoids getting trapped in local minima and provides the parameter set that globally gives the best fit to the data.

First, we tried fitting a simpler model than the independent-donor model described above to the empirical data shown in Figure 1B&D. This simpler model did not contain time delays (*τ*_*d*_, *τ*_*l*_, *τ*_*p*_), differential infection parameters (*a*_*p*_, *p*_*l*_) or parameter dependence on bacterial and phage densities (in *α, p* and *l*). Hence, only conjugation (*p*), adsorption (*α*), induction (*i*) and lysogenization (*l*) rates were fitted, which however did not result in a good qualitative fit (Figure S2).

Accordingly, we included several, biologically more realistic features into the model (see independent-donor model description) as our empirical data suggested that time delays, phage density-dependence in lysogenization probability as well as bacterial density-dependence in plasmid conjugation and phage infection were relevant in our system (Figure 1B). For plasmid conjugation, we ended up fitting a conjugation rate (*p*), its growth dependence (*k*_*p*_) and a time delay (*τ*_*p*_) to the single-donor conjugation experiment in Figure 1B. As phage infections are more complex, the empirical data from the single-donor lysogenization experiment was not sufficient to fit all of the new phage infection parameters. Hence, we used literature values as a starting point (see Table S2 and fitted adsorption rate (*α*) in LB and SM, as well as its growth dependence (*k*_*a*_, *k*_2_), induction rate (*i*) and the lysogenization factor (*l*^*′*^) to data from Figure 1D& S1.

The surprisingly low number of double-transformants in the independent-donor experiments in contrast to their high number in common-donor experiments suggested that infection rates for MGE-carrying cells might not be the same as for MGE-free cells. Keeping the other parameters constant, we fitted scaling factors for conjugation to lysogens (*p*_*l*_) and adsorption to plasmid-carriers (*a*_*p*_), to the independent- and common-donor experiments individually. For the independent-donor experiments, the best fit showed a 10-fold decrease in conjugation to lysogen cells, but no effect of plasmid-carriage on phage infection. For the common-donor experiments however, fitting showed that both rates should be increased, adsorption to plasmid-carriers about 10-fold and conjugation to lysogens even 500-fold in order to explain the increase in double carriers over lysogens (3B,C). The parameter values used are summarized in Table S1&S2 for the independent-donor model and in Table S3&S4 for the common-donor model.

#### Literature values

The remaining phage parameters, which were likely to be similar to previous measurements, were taken from the extensive empirical literature on phage *λ*: burst size (*β*) [28], the time delay in phage decision (*τ*_*d*_) [30] and the phage decay rate (*γ*_*V*_) [1, 6].

##### Model comparison

In order to test the importance of the various potentially biologically-relevant additions to the model, we compared the quality of the model fit between our ‘complex’ model and simpler versions using AIC (Akaike information criterion) (Figure S2A). Specifically, we compared the ‘full’ model to i) one without time delays, ii) one without bacterial density dependence, iii) one without differential infection, iv) one without i)–iii) and v) the simple model from above, where, additionally to iv) lysogenization rate is not phage density dependent. We calculated the residual sum of squares (RSS) between the model simulations and empirical data for each of the replicates in the single-donor, independent-donor, common-donor and MGE-transfer control experiments (Figure 1B,D,3B,C,S1) and summed these errors. The difference in AIC between model versions was then calculated by correcting for the number of parameters (*k*_*par*_) used in the models and the number of observation points (*n*_*obs*_) used to compare model simulations and empirical data [31]:

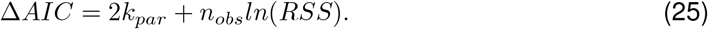

(Δ*AIC* differs from absolute AIC by an additive constant, which is the same for all model variants, as it is only dependent upon the data points, and does not contribute to differences in goodness of fit [31].) This approach allowed us to identify differential infection as a crucial component of our model fit.

The additional influence of time- and density-dependence in the model parameters underlines the highly dynamic nature of plasmid and phage infections and the importance of capturing it.

##### Parameter sensitivity analysis

In order to explore the global parameter sensitivity of our model, we used (uniform) random sampling of parameters within biologically realistic ranges (Table S2) to obtain 1000 random parameter combinations. We recorded the number of transconjugants, newly formed lysogens and double-transformants after 24h and visualized their spread across all 1000 different parameter sets using violin plots (Figure S2B,C). We compared this spread between random parameter sampling done for the ‘simple’ and the ‘full’ model version and found little difference in the variation of the output. Generally, tranconjugants and lysogens are present at high numbers (≥10^5^), whereas the number of double-transformants are more variable, as they compound variations in the number of transconjugants and lysogens (Figure S2B,C).

Further, we explored the impact of several infection parameters on bacterial population numbers in more detail for the independent-donor model by varying adsorption rate [32, 33], conjugation rate [34], lysogenization probability [35], induction rate [23, 24, 35, 36], burst size [37], growth rate, and starting bacterial density over a wide range of parameters, chosen based on empirically observed values. In simulations where parameters other than adsorption rate were varied, we used the lower adsorption rate of LB environments, though we verified that using the higher adsorption rate of SM environments did not change our results notably. Contour plots show the absolute cell numbers of various subpopulations at the end of a 24h-simulation run.

##### Common-donor model

(Figure 3D, Table S3& S4)

The common-donor model is a simplified version of the independent-donor model, starting from a recipient (*R*) and a common donor, *D*_*LP*_, which contains the plasmid and the prophage. Hence, the donor can transfer phages and plasmids (individually), yielding recipient plasmid carriers (*R*_*P*_):

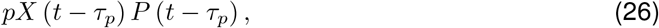

with *X* ∈ {*R, R*_*L*_}, and *P* = *R*_*P*_ + *D*_*LP*_ + *R*_*LP*_ ; as well as recipient lysogens (*R*_*L*_):

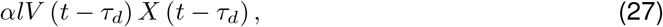

for *X* ∈ {*R, R*_*P*_}, after the respective time delays. (As the donor already carries both MGEs, there is no transfer to the donor subpopulation, as opposed to the independent-donor model.) Analogous to the independent-donor model, phages can infect cells either productively (lytic pathway) or integrate into the host genome (as prophages), which are induced at a low spontaneous rate *i*. Phage adsorption and plasmid conjugation are dependent on host density, while lysogenization rate is dependent on free phage virion density as described in Equations 20-22 of the independent-donor model.

Further infection of recipients carrying already one MGE (*R*_*P*_ or *R*_*L*_) produces double-transformants (*R*_*LP*_). Fitting of simulations to empirical data showed that phages preferentially adsorb to plasmid carriers at rate

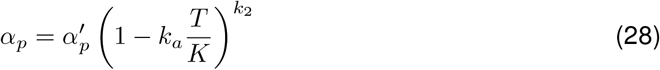

and plasmids conjugate preferentially to lysogens at rate *p*_*l*_ (Equation 22), as compared to MGE-free recipients (see ‘Parameter estimation from empirical data’).

## Results

We investigated the transfer dynamics of a conjugative plasmid in the presence of temperate phages by combining experiments of plasmid spread in the absence (single-donor experiment) or in the presence of a prophage-carrying population (independent-donor and common-donor experiment) with mathematical modelling. Experiments were performed with phage *λ* and plasmid RP4, as they are both well-studied systems, which, to our knowledge, do not contain systems affecting the other MGE’s transfer. We used a ‘standard’ liquid environment (rich medium, ‘LB’), allowing efficient transfer of both MGEs, as well as a ‘ phage-friendly’ environment (LB supplemented with SM-buffer, ‘SM’), which benefits phage infections by increasing adsorption rates [38].

### Plasmid spread starts later than prophage spread, but continues for longer

First, we compared transfer dynamics of temperate phages (lysogenization) and plasmids (conjugation) independent of the respective other MGE (Figure 1A). In both environments, the mean number of newly formed lysogens increased considerably earlier than the formation of transconjugants (Figure 1B). Since prophage spread saturated earlier, transconjugants reached a higher number than lysogens after 24 hours in LB (Figure 1B). In SM, the number of lysogens over time was significantly higher compared to LB (linear model at T24: *F*_1,4_ = 13.09, *p* = 0.022). This explains the approximately equal numbers of transconjugants and lysogens in SM after 24 hours, despite the earlier saturation of prophage spread. Hence, the high concentration of Mg^2+^ and Cl^-^ ions in the SM environment favoured phage adsorption, but did not affect conjugation (linear model at T24: *F*_1,6_ = 1.09, *p* = 0.34; Figure 1B).

### Phage infections limit plasmid spread in environments with two independent donors

Next, we determined the formation of plasmid- and prophage-carrying cells in co-culture with two independent donors (one plasmid donor and one phage donor) and one recipient (Figure 1C). In agreement with the single-donor experiments, lysogens emerged faster (i.e., recipients and plasmid donors that acquired the prophage) than transconjugants. However, within the given timeframe of 24 hours, the total amount of transconjugants was not able to exceed the amount of lysogens. Further, the final amount of transconjugants was almost two orders of magnitude lower in LB and SM after 24 hours as compared to the single-donor experiment (linear model: T24 LB: *t*_17_ = 5.049, *p <* 0.001; T24 SM: *t*_17_ = 9.61, *p <* 0.001). This indicates a strong negative impact of temperate phages on plasmid transfer. In contrast, phage transfer was not negatively affected by plasmids. Here, the initial increase in lysogens was even higher in SM compared to the single-donor experiment (linear model: T3 LB: *t*_11_ = −5.174, *p* = 0.002; T5 SM: *t*_12_ = −8.82, *p <* 0.001). However, in the single-donor experiment, the starting ratio between phage donors and recipients was 1:1, whereas in the independent-donor experiment phage-susceptible bacteria (recipients and plasmid donors) outnumbered phage donors 2:1. This allowed for more phage amplification and presumably resulted in a faster increase of lysogens in the independent-donor experiment.

### A mathematical model identifies crucial infection parameter dependencies

While it is intuitive that phages limit or slow down plasmid spread through bacterial killing, we also observed a low abundance of double-transformants (recipients that acquired the plasmid and the prophage) throughout the experiment. This was surprising given the high numbers of transconjugants and newly formed lysogens after 5h (Figure 1D). We thus used a mathematical model to explore this puzzle and to identify infection processes that determine the numbers of transconjugants, lysogens and double-transformants.

For the fitting of model parameters (Methods, Table S2), we took advantage of our empirical data on growth rates (Table S5) and transfer dynamics between various cell types (Figure 1B, S1) as well as the available literature. A simple model assuming constant and instantaneous phage and plasmid transfer however did not yield a good qualitative fit to empirically observed dynamics (Figure S2). Accordingly, we tried to match realistic infection dynamics more closely by considering time delays associated with the process of plasmid conjugation and the time needed for reproduction inside the cell during lytic phage cycles [39]. Further, our data, and other empirical studies [40, 41] show decreased plasmid and phage spread on stationary phase bacteria, indicating that adsorption and conjugation are dependent on bacterial growth rate. Lysogenization rate on the other hand increases with phage density [42].

Allowing additionally for differences in conjugation and adsorption to MGE-free and MGE-carrying cells gave the best fit with a 10-fold reduced plasmid transfer to lysogens (but no change in adsorption due to plasmid carriage) (Figure S2). In combination with phage-mediated killing of plasmid-carrying cells, this reduced plasmid transfer could explain the decrease in transconjugants compared to the single-donor and transfer control experiments (Figure 1B,D, S1), as well as the low number of double-transformants. Altogether, incorporating time delays, density-dependent infection parameters and differential infection resulted in a much better quantitative fit to the observed trajectories of lysogens, transconjugants and double-transformants (Figure 1, S2, S3). Individually, time delays and density-dependence only showed a small impact on the fit, whereas differential infection was quite influential on its own (Figure S2). Thus, even in this deceptively simple system, consisting of only three players – temperate phages, plasmids and bacterial hosts – the interactions are highly dynamic and complex.

### Transconjugant formation is dependent on the conjugation rate, but largely independent of phage infection parameters

Given the complexity of the interactions in our system demonstrated by the model, and based on its good quantitative fit with empirical observations, we used the model to explore different plasmid-phage combinations by varying infection parameters over a wide range that captures empirical values (Methods). We focused on phage adsorption and plasmid conjugation rate, which, in this system, are the most important fitness parameters for horizontal phage and plasmid transfer. In addition, we also explored how other phage infection (lysogenization probability, induction rate, burst size), as well as host cell (growth rate, starting density) parameters influence the numbers of the individual subpopulations.

Contrary to our expectations, we found that the number of transconjugants after 24h was largely independent of the adsorption rate, and almost exclusively depended on the conjugation rate (Figure 2A, S4A): a high number of transconjugants could always be obtained at high conjugation rates (optimally below 10^−10^, which is well within empirically observed values [43]). Similarly, the number of newly formed lysogens was primarily determined by the adsorption rate, varying substantially across empirically measured values [32], with our fitted values for SM being closer to the higher end (Figure 2B, S4B). In contrast, the emergence of double-transformants was limited by conjugation and adsorption rate, which narrowed down the window in which high double-transformant numbers can be expected (Figure 2C, S4). Our model indicated that the number of double-transformants could be substantially increased through higher conjugation as well as adsorption rates (Figure 2A, C).

**Figure 2:**
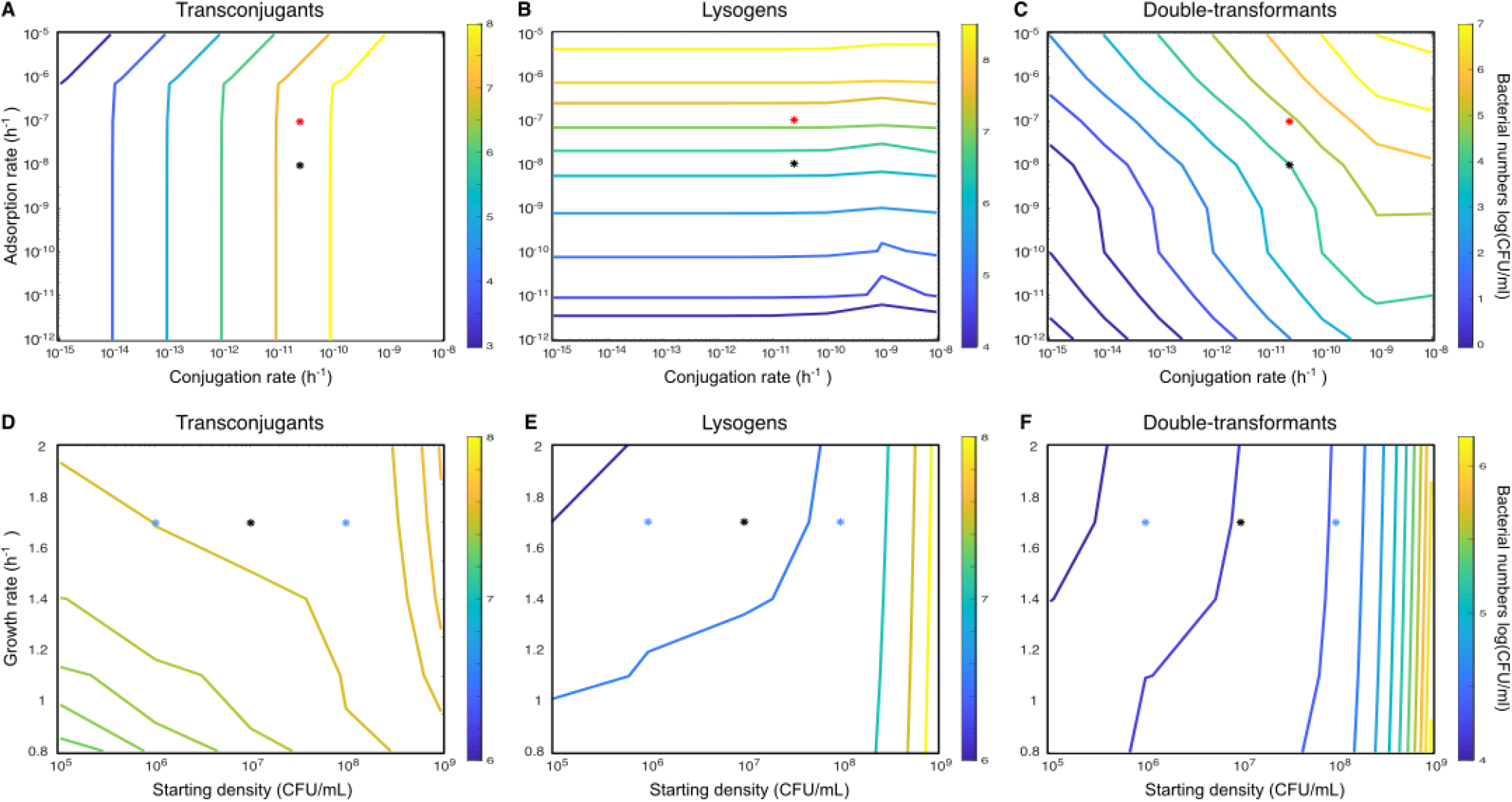
Variation of modelling parameters using the independent-donor model. Contour plots give bacterial cell numbers (log(CFU/mL)) of (A, D) transconjugants, (B, E) new lysogens and (C, F) double-transformants in response to variation in adsorption and conjugation rate (top row), or growth rate and starting bacterial density (bottom row) at the end of 24h-simulations. D-F were simulated using parameters from LB environments. Asterisks show the parameter values used in the simulations for Figure 1F in LB (black) or SM (red) or for Figure 3E&F (blue).

When we varied the lysogenization rate, we observed the highest numbers of newly formed lysogens and double-transformants at lower values (Figure S5), because a high probability of lysogenization resulted in low numbers of free phages for new infections. Similarly, higher induction rates produced a higher number of free phages and led to increased lysogen formation (Figure S5). Burst size only affected the time at which phage infection takes off, but not the final number of lysogens (Figure S5). Lysogenization probability, induction rate and burst size all showed almost no effect on transconjugant numbers and a relatively weak effect on double-transformant numbers (Figure S5).

The influence of host parameters, i.e., growth rate and starting density, showed a weak influence on subpopulation numbers overall (Figure 2D-F), indicating that density-dependence affects the infection dynamics more than the final numbers. Higher starting densities slightly increased the number of transconjugants, lysogens and double-transformants. Note, that while this is due to the mass-action principle for conjugation, lysogenization is increased because we assume that a higher number of starting lysogens means a higher number of free (spontaneously induced) phages at the beginning of the experiment (see Methods: ‘Initial conditions’) – which in turn increases lysogenization frequency. Assuming no free phages in the beginning would lead to a decrease in lysogen numbers with starting density as stationary phase would be reached before the infection can take off.

### Transconjugants gain an advantage in common-donor experiments

According to the model, transconjugants and particularly double-transformants should increase with higher conjugation rates (Figure 2A-C). As this is difficult to achieve experimentally without changing many other factors, we decided to favour plasmid spread in a different manner, namely by using a common donor, carrying both, the phage and the plasmid (Figure 3A). This avoids killing of plasmid donors due to lytic phage infections – as observed in the independent-donor experiments (Figure S3A). Further, we experimentally tested the prediction that MGE transfer will be largely independent of bacterial starting densities (Figure 2D-F).

**Figure 3:**
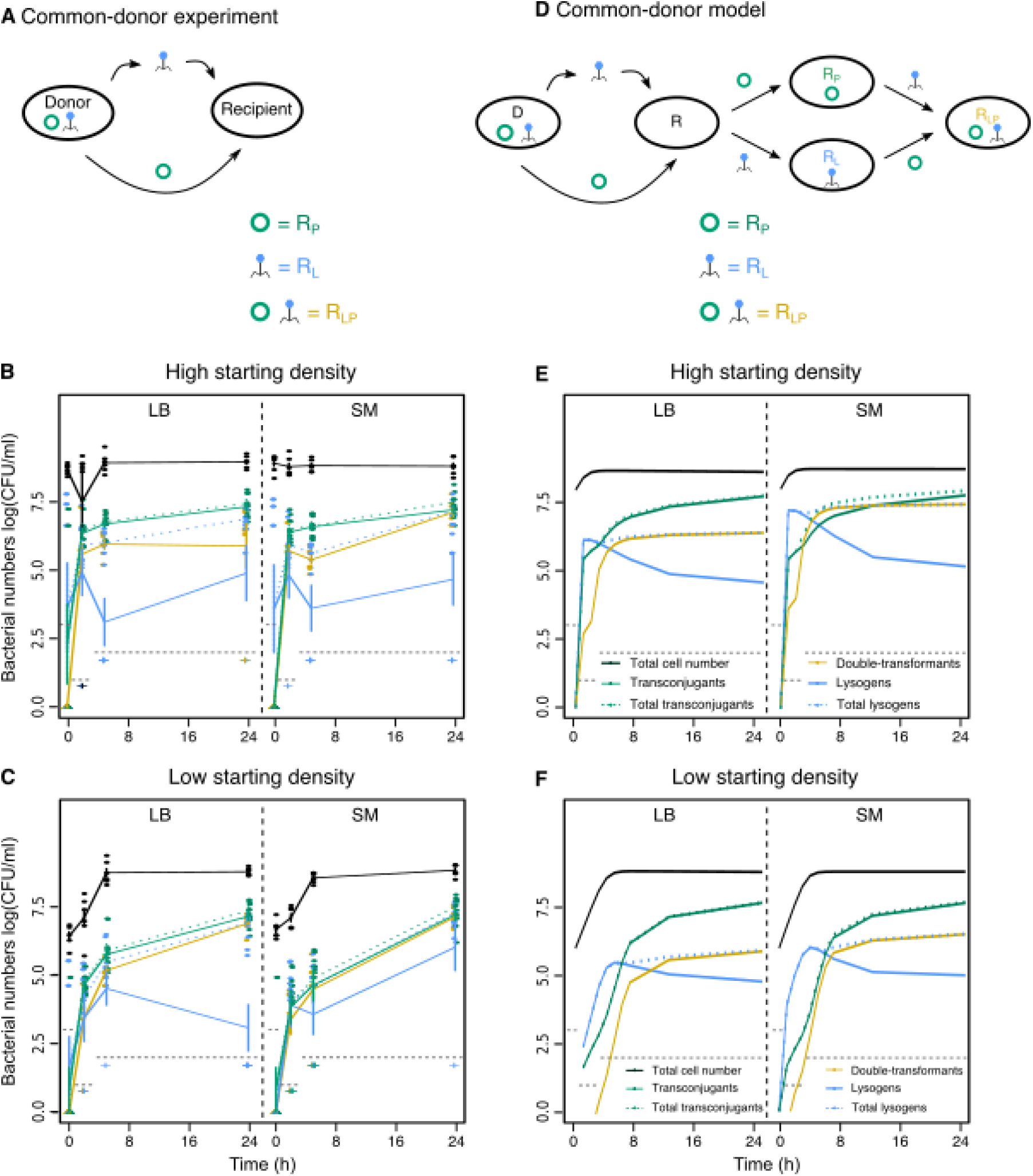
Common-donor experiment. (A) A common donor carrying the plasmid and the phage (*D*_*LP*_) was mixed with recipient cells. Transconjugants (green) are recipients, which acquired the plasmid (*R*_*P*_), lysogens (blue) are recipients, which acquired the prophage (*R*_*L*_), and double-transformants (yellow) are recipients, which acquired the plasmid and the phage (*R*_*LP*_). (B, C) Total cell number (black), as well as newly formed lysogens (blue), transconjugants (green), and double-transformants (yellow) measured as colony forming units log(CFU/mL) over time in LB (left panel) or SM (right panel) starting with (B) high or (C) low bacterial densities (shown are single data points as well as mean ± s.e., n=6). For comparison with Figure 1B total transconjugants (*R*_*P*_ + *R*_*LP*_) and total newly formed lysogens (*R*_*L*_ + *R*_*LP*_) are shown in dashed green or blue lines but might be hidden by the respective solid lines. Grey dashed lines indicate the empirical detection limit at a given time point (Methods). (D) Schematic of reactions for the common-donor model. (E, F) Predictions of 24h-model simulations for phage and plasmid spread with a common donor in LB (left panel) or SM (right panel) starting with (E) high or (F) low bacterial densities using highly preferential infection parameters of MGE-carriers (colours and line types as described in B&C).

As opposed to the independent-donor experiments, transconjugants appeared initially at higher numbers than lysogens in the common-donor experiments. The faster initial rise of transconjugants for low density treatments was puzzling (Figure 3B, C), given the lag of transconjugant appearance in single-donor experiments (Figure 1B). As predicted, double-transformants seemed to reach higher absolute numbers in both environments as compared to the independent-donor experiments, which was – potentially due to the empirical detection limit, which reduced the statistical power – not significant (linear model: T24 SM: *t*_10_ = −1.87, *p* = 0.09, T24 LB: *t*_10_ = −2.16, *p* = 0.056). The number of total lysogens (i.e., newly formed lysogens and double-transformants) was however significantly lower in SM as compared to single- and independent-donor experiments (linear model, T24 SM: single-donor experiment: *t*_16_ = 4.9, *p* = 0.0012, independent donor: *t*_16_ = 5.62, *p* = 0.0003; Figure 1, 3), and the number of cells carrying only the prophage was very low overall as they were quickly turned into double-transformants. In contrast to the independent- (linear model: T24: *F*_1,7_ = 40.11, *p <* 0.001) and single-donor (linear model: T24: *F*_1,4_ = 13.09, *p* = 0.022) experiments, phage adsorption (LB vs. SM) did not influence the total number of lysogens in the common-donor experiment (linear model: T24 high density: *F*_1,10_ = 2.02, *p* = 0.186; T24 low density: *F*_1,10_ = 4.61, *p* = 0.057).

In agreement with our predictions, the dynamics were independent of bacterial starting densities. Using the parameters determined from the independent-donor experiments however poorly predicted the number of double-transformants and lysogens in the common-donor experiments (Figure S6). We could substantially improve the model fit (Figure 3, S2A) when we allowed highly preferential infection of recipients already carrying the other MGE (10-fold increased adsorption to plasmid carriers and 500-fold increased conjugation to lysogens, Table S4) – as opposed to the lower conjugation rate to lysogens in the independent-donor experiment (Table S2). Using the infection parameters fitted to the common-donor experiments would increase the absolute number of transconjugants and double-transformants substantially in the independent-donor simulations (Figure S7), contrasting with our empirical observations (Figure 1D). This indicates that there are unknown, but biologically relevant interaction mechanisms between the plasmid and the phage in our system, and that infection parameters related to ‘co-infection’ of a cell can depend on the genetic background of the donor and the recipient.

## Discussion

Here, we explored to which extent the presence of active prophages in a host population can limit conjugative plasmid transfer. While we observed the intuitive effect of all phages – killing of bacterial cells – we also reveal that understanding the magnitude and temporal dynamics of this death factor in different environments becomes more complex in the presence of temperate phages.

Already the comparison between plasmid and phage transfer in single-donor experiments revealed interesting differences in their transfer dynamics (Figure 1B): while lysogenization occurred rapidly after mixing with recipient cells, plasmid conjugation lagged behind by several hours. This might be surprising, given that for phage infection and subsequent lysogenization to occur at significant frequencies, prophages have to be induced (which only happens at a low spontaneous rate) and a certain phage-to-bacteria ratio has to be reached [14–16]. However, we likely carried over a number of free phages from the diluted overnight lysogen cultures, which we calculated to be around two orders of magnitude lower than the respective lysogen starting density (Methods). Further, in agreement with our data, it has been shown before that conjugation frequency for some plasmids increases only slowly over the exponential growth phase and peaks at the transition to stationary phase [44]. Another, mutually non-exclusive, explanation could be that recipients pay an acquisition cost (in addition to the overall plasmid metabolic burden), which slows down initial growth and spread of transconjugants [45]. Even though transconjugant formation started later, it continued longer, allowing conjugation even after the bacterial population reached stationary phase.

Our mathematical model showed that these unequal transfer dependencies on host cell growth, as well as the duration of the infection process (which is also dependent on host cell growth), are relevant in reproducing empirical observations. Hence, plasmid and phage spread in a common environment strongly depends on their respective host exploitation strategies (e.g., horizontal transfer in exponential versus stationary phase bacteria). The ability of some plasmid types to conjugate at normal or even higher rates during stationary phase [46] could be an attempt at ‘catching up’ when most temperate (and many lytic) phages are unable to propagate. While the influence of host cell physiology has been studied for each system independently [27, 44], its impact on concomitant infection dynamics of phages and plasmids is hard to predict and will require further experiments.

With temperate phages, the potential for killing surrounding cells depends not only on bacterial numbers, but also on phage density, often producing only a relatively short burst of (notable) lytic activity [4]. According to our independent-donor experiments, this short burst is enough to substantially limit the total number of transconjugants (Figure 1D). Transconjugant dynamics in the presence of lysogens can be approximated by interpreting phages as an additional ‘death rate’ for non-lysogen cells (plasmid-donors, recipients and recipient-transconjugants, Figure S8). As prophages are pervasive in natural isolates [4], these findings are highly relevant for understanding the spread of plasmids or other MGEs in natural environments. Further, the presence of prophages in conjugation assays could distort experimental measurements of conjugation rates, potentially leading to environment- or growth rate-dependencies caused by other factors than plasmid transfer.

Temperate phage killing or protection of cells, which phages, but also plasmids rely on for survival, introduces a feedback into the system, which results in a complex dependence of the phage-mediated ‘death rate’ on environmental conditions. This complexity becomes visible through the significantly higher number of (total) transconjugants achieved after 24h, when plasmids were allowed to spread from a common instead of an independent donor (Figure 1D, 3B, C). Moreover, transconjugant formation also started earlier when plasmids spread from a common donor (Figure 3). This effect might be caused by induction of lysogens continuously killing a small part of the plasmid phage donor population, meaning that overnight cultures might undergo more turnover [47], and that a larger fraction of the population might still be capable of conjugation [44]. The faster initial plasmid spread and the reduced killing of plasmid donors (which is now restricted to spontaneous induction) in common-donor experiments effectively increased plasmid conjugation rate, which according to our model allowed overcoming the ‘ phage-mediated ‘death rate’ (Figure 2A, 3E, F). Indeed, our empirical data showed higher transconjugant and particularly double-transformant numbers in the common-donor experiments. The result that higher conjugation rates can overcome the inhibition of plasmid spread through phage killing was largely independent of phage infection parameters, and hence should be valid for any particular prophage type. Accordingly, we would expect a higher selection pressure on plasmids to evolve high conjugation rates in prophage-rich environments. This might however be detrimental for other reasons, like metabolic burden on host cells, and would indicate that the frequent occurrence of phage-inhibitory systems on conjugative plasmids [5, 6, 9] serves not just as a defence against lytic phages but also against temperate phages. In some cases, plasmid entry into a host cell was even observed to result in curing of resident prophages [48].

Our study suggests that the speed of transfer depends on the MGE-environment combination. Not only plasmids, but also temperate phages can contribute to horizontal gene transfer, using a plethora of mechanisms, which include specialized and generalized transduction (packaging of non-phage genes) or phenotypic changes through the expression of prophage-encoded genes (lysogenic conversion) [49]. In our independent-donor experiment, conjugation seemed to be the limiting factor for the number of transconjugants and double-transformants. Hence, we expect that in environments or plasmid types with low conjugation rates, prophages would be more efficient resistance or virulence gene spreaders (Figure 2A-C). Similarly, in environments with prophage types characterized by low adsorption rates we expect plasmids to be the dominant spreader of resistance or virulence genes. When conjugation and adsorption rate both are relatively high, double-transformants will be abundant and horizontal gene transfer will be mediated efficiently by both, plasmids and temperate phages. As it is beneficial for accessory genes, capable of increasing bacterial fitness in a specific environment, to be transferred by the preferred mode in that environment, interactions between prophages and plasmids could be instrumental in determining the location of horizontally transferred genes.

While our model generally showed a good quantitative fit, the difficulty in reproducing the abundance of double-transformants – which our simulations slightly overestimated in the independent-donor experiment (Figure 1D, F) and underestimated in the common-donor experiment (Figure 3) – indicates that the biological processes underlying infection of and from an MGE-carrying cell are more complex than those of wildtype cells. Interestingly, the model fit was strongly improved by assuming that prophages inhibited plasmid entry to some extent, if plasmids and phages come from independent donors, and conversely by assuming preferential infection of cells carrying the other MGE for common-donor experiments (Figure S6). This is particularly noteworthy, as time delays and density-dependence played a comparatively minor role to differential infection in improving the fit with empirical observations (Figure S2). Our common-donor results demonstrate that, for plasmids, it could be beneficial to preferentially infect lysogens as co-resident prophages will confer superinfection immunity. However, for prophages, co-existence with a plasmid in a common donor did not increase lysogen formation, which could lead to an evolutionary conflict between prophages and conjugative plasmids.

Facilitation of co-infection by another MGE could for instance stem from mechanisms that disable host cell defences, such as anti-restriction-modification systems, which have been found on both, plasmids [4] and phages [50]. The RP4 plasmid as well as the *λ* prophage used in this study encode anti-restriction systems, which are active against the EcoKI restriction enzyme in *E. coli* K12 [50, 51], and could potentially facilitate infection of a cell already carrying the other MGE. Intriguingly, fitting of the model parameters to empirical data indicated that interactions between plasmids and prophages were reversed when adapted to sharing of the same host cell. This is difficult to explain via anti-restriction systems alone as they should not lead to antagonistic effects with independent-donors, and there are, according to our knowledge, no other obvious candidate genes for this kind of interaction found on RP4 or *λ*. Hence, especially for prophage spread, future research needs to tease apart the individual processes during concomitant or sequential infection and include more mechanistic realism into the model in order to identify relevant interactions.

In summary, we find that the presence of active prophages in the environment limits plasmid transfer by introducing an additional death rate, the magnitude of which is determined by the specific population dynamics and non-trivial to predict for different environmental conditions. Notably, either prophages or plasmids can increase their own transfer largely independently of the specific plasmid-prophage pair. Further, the dependence of transfer efficiency on host physiology could also impact the effect of lytic phages on plasmid spread. These results open up new avenues of exploration regarding phage-plasmid interactions and set up expectations for plasmid evolution in nature. According to our model, we predict that the evolution of higher conjugation rates, as well as co-infection of host cells carrying prophages, is beneficial for conjugative plasmids in environments where prophages are abundant.

## Acknowledgements

We thank A. Hall for his valuable input during experimental design and data interpretation, and T. Roth for his support during the experimental work. Special thanks go to H. Chabas for her comments on an earlier version of this manuscript and to R. Regös for helpful discussions and comments. This project has received funding from the Swiss National Science Foundation (grant number PZ00P3_179743) given to CCW and was supported by an ETH Zurich Postdoctoral Fellowship (19-2 FEL-74) received by CI.

