## Supplementary material for "Conjugative plasmid transfer is limited by prophages but can be overcome by high conjugation rates"

### Supplementary Figures

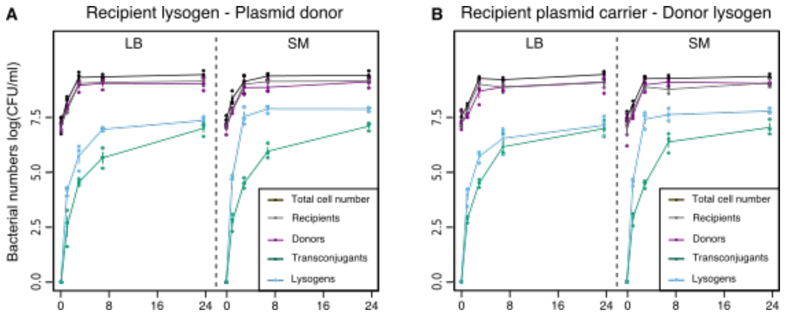


Figure S1: **Transfer rates to cells already carrying an MGE (MGE-transfer control experiments).** Total amount of bacterial cells (black), recipients (grey), donors (purple), newly formed transconjugants ($R_{LP}$ in A, $D_{LP}$ in B; green) and newly formed lysogens ($D_{PL}$ in A, $R_{PL}$ in B; blue) measured as colony forming units log(CFU/mL) over time in LB (left panel) or SM (right panel) is shown for MGE-transfer control experiments between (A) recipient lysogens ($R_L$) and plasmid donors ($D_P$), or (B) recipient plasmid carriers ($R_P$) and donor lysogens ($D_L$). Note that killing of plasmid carriers by infecting phages is hard to determine from comparison with lysogens as lysogens also get killed to some extent by spontaneous induction. Shown are individual data points as well as mean ± s.e., n=3.

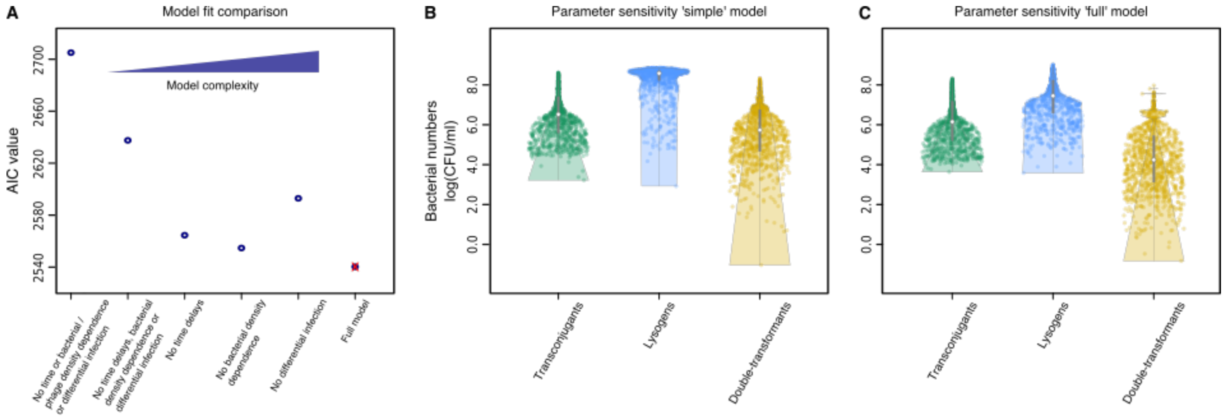


Figure S2: **Empirical fit of different model variations and global parameter sensitivity analysis.** (A) AIC values are shown for model variants of increasing complexity, calculated from the model fit to experimental data of single-donor, independent-donor, common-donor and transfer control experiments. The 'full' model used throughout the manuscript is indicated by a red cross. (B, C) Violin plots show the model output for 1000 randomly chosen parameter combinations using uniform sampling within biologically realistic ranges (Table S2). Bacterial numbers of newly formed transconjugants (green), lysogens (blue) and double-transformants (yellow) after 24h are shown as individual dots for each parameter combination, either using the (B) 'simple' (leftmost model in A) or (C) 'full' model version (rightmost model in A).

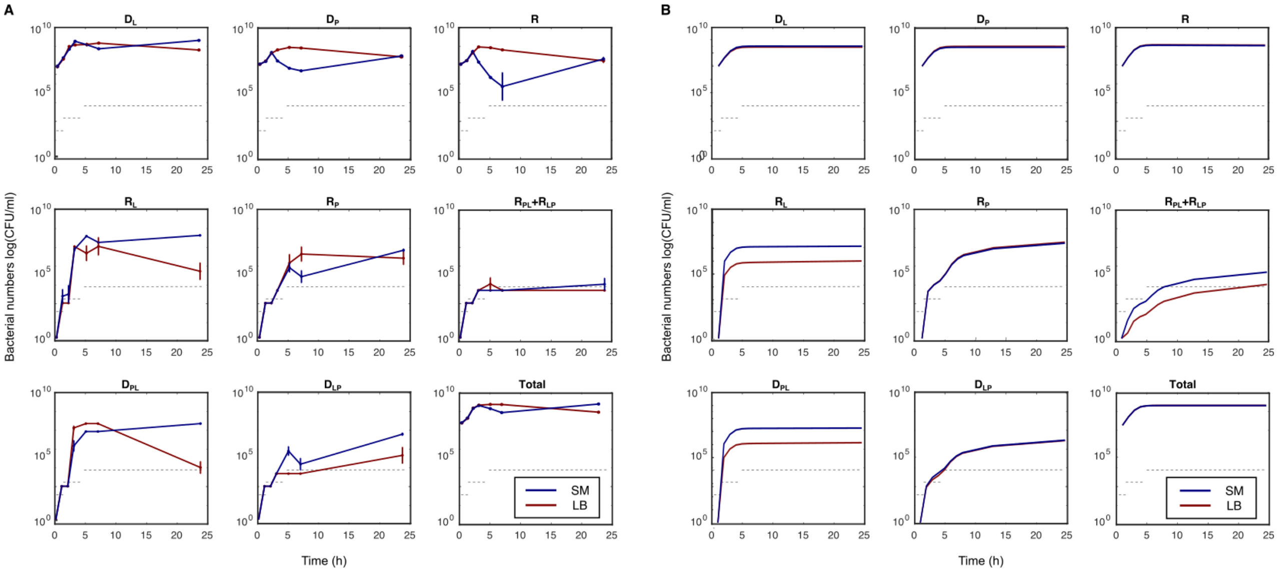

Figure S3: **Model fit to independent-donor subpopulation data.** Bacterial cell numbers log(CFU/mL) are shown for (A) empirical data (mean $\pm$ s.e., n=6) and (B) model simulations over 24h in LB (red) or SM (blue) for phage donors ($D_L$), plasmid donors ($D_P$), recipients ($R$), recipient lysogens ($R_L$), recipient plasmid carriers ($R_P$), double-transformants ($R_{LP} + R_{PL}$), plasmid donors with lysogens ($D_{PL}$), phage donors with plasmids ($D_{LP}$) and total cell numbers. Grey dashed lines indicate the empirical detection limit at the given time point.

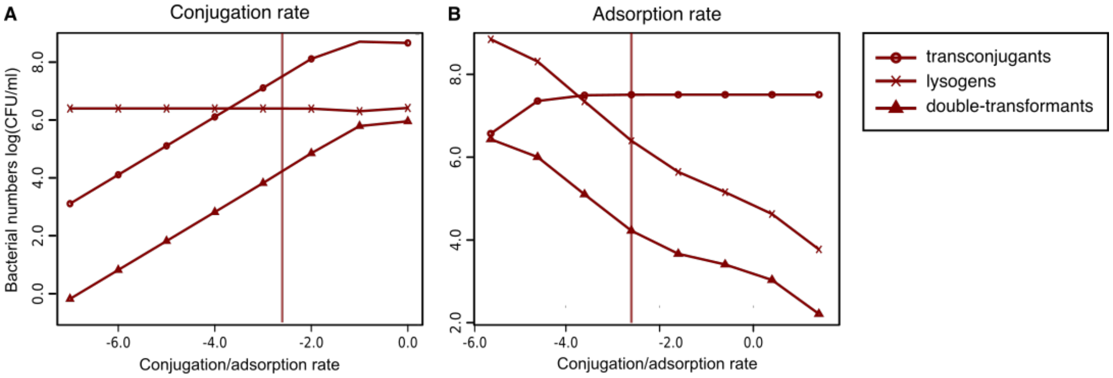


Figure S4: **Variation of conjugation and adsorption rate using the independent-donor model.** Bacterial numbers (log(CFU/mL)) are shown when (A) plasmid conjugation rate or (B) phage adsorption rate were varied over a range of parameters. The x-axis gives the ratio of conjugation to adsorption rate. Shown are changes in the number of transconjugants (circles), lysogens (crosses) and double-transformants (triangles) at the end of 24h-simulations in LB. Vertical lines show the (ratio of the) parameter values used in the simulations for Figure 1F.

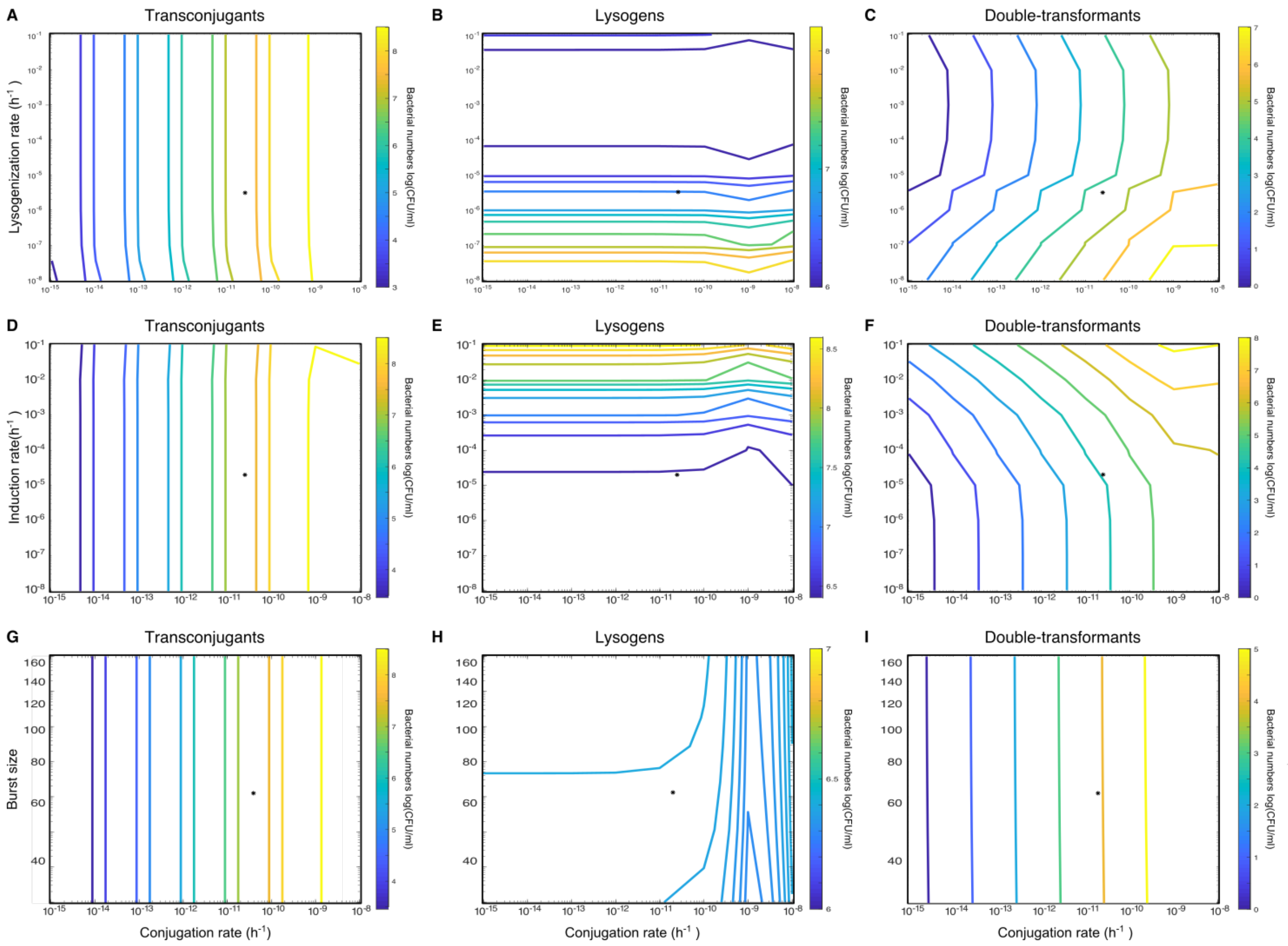


Figure S5: **Variation of the scale parameter for the lysogenization rate, induction rate or burst size together with the conjugation rate using the independent-donor model.** Contour plots give the cell number (log(CFU/mL)) of (A,D,G) transconjugants, (B,E,H) newly formed lysogens and (C,F,I) double-transformants at the end of 24h-simulations in LB. Stars show the parameter values used in the simulations for Figure 1F. Contrary to intuition, the number of lysogens decreases with higher lysogenization rate, as increased lysogen formation at the beginning of the infection would inhibit the production of free phages. Neither of the three parameters showed a strong influence on the number of double-transformants (as compared to conjugation rate.)

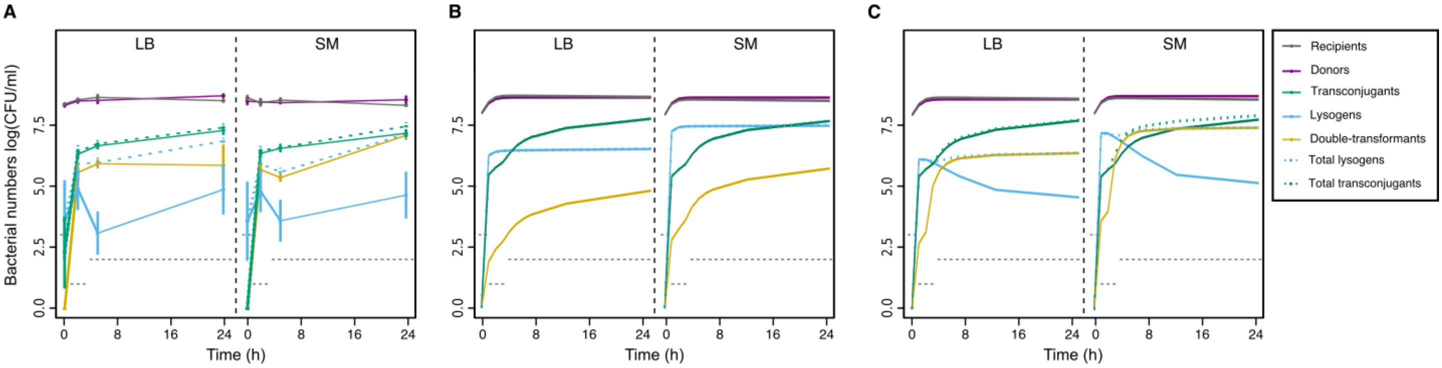


Figure S6: **Model fit to common-donor subpopulation data.** Bacterial cell numbers (log(CFU/mL)) of recipients (grey), plasmid phage donors (purple), transconjugants (green), newly formed lysogens (blue), double-transformants (yellow), total transconjugants (i.e., including double-transformants; dashed green) and total lysogens (i.e., including double-transformants; dashed blue) are shown for (A) empirical data (mean ± s.e., n=6) and (B,C) model simulations over 24h in LB (left panel) or SM (right panel) starting from high population densities. Simulations were done either using parameters from the (B) independent-donor fit, or (C) using preferential infection of MGE-carrying subpopulations. Note that the dashed lines of total MGE-carriers are sometimes hidden beneath the corresponding solid lines. Grey dashed lines indicate the empirical detection limit at the given time point (dynamics for low population densities look similar and are not shown).

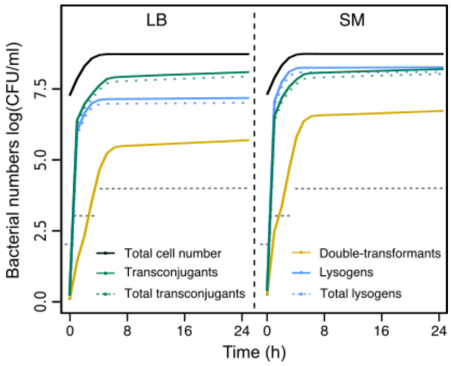


Figure S7: **Model predictions for the independent-donor experiment with the preferential infection fit from the common-donor model.** Predictions of 24h-simulations of bacterial cell numbers in log(CFU/mL) for total population size (black), transconjugants (green), newly formed lysogens (blue), double-transformants (yellow), total transconjugants (i.e., including double-transformants; dashed green) and total lysogens (i.e., including double-transformants; dashed blue) in LB (left panel) or SM (right panel) using parameters fitted to the common-donor model (Figure 3E&F). Note that the dashed lines of total MGE-carriers are sometimes hidden beneath the corresponding solid lines. Grey dashed lines indicate the empirical detection limit at the given time point. The number of transconjugants and double- transformants is much higher than experimentally observed (Figure 1D).

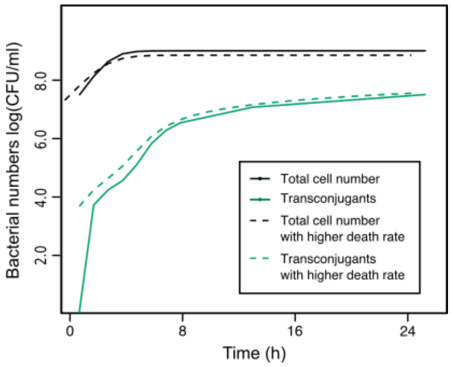


Figure S8: **Phages as death rates.** 24h-simulations in LB of bacterial cell numbers in log(CFU/mL) for total population size ($R + R_P + D_P$; black) and total transconjugants (green) from simulations including phage infections are shown with solid lines (dashed line in Figure 1F for total transconjugants), and from simulations with an additional death rate for recipient and plasmid-carrying cells of $0.5\text{h}^{-1}$ are shown with dashed lines.

### Supplementary Tables

Table S1: Model variables used in the independent-donor model.

| Model variables | Initial values (SM/LB) | Parameter range | Description | Genetic background |
|---|---|---|---|---|
| $D_L$ | $10^7$ cells | $10^5 - 10^9$ cells | Phage donor | Prophage |
| $D_P$ | $10^7$ cells | $10^5 - 10^9$ cells | Plasmid donor | Plasmid |
| $R$ | $10^7$ cells | $10^5 - 10^9$ cells | Recipient cells | MGE-free |
| $R_L$ | 0 | - | Recipient lysogens | Prophage |
| $R_P$ | 0 | - | Recipient plasmid cells | Plasmid |
| $D_{LP}$ | 0 | - | Phage donor plasmid cells | Prophage and plasmid |
| $D_{PL}$ | 0 | - | Plasmid donor lysogens | Prophage and plasmid |
| $R_{LP}$ | 0 | - | Recipient lysogens with plasmid | Prophage and plasmid |
| $R_{PL}$ | 0 | - | Recipient plasmid lysogens | Prophage and plasmid |
| $V$ | $5 \cdot 10^5$ virions | $5 \cdot 10^3 - 5 \cdot 10^7$ virions | Free phage virions | - |
| $T$ | $3 \cdot 10^7$ cells | $3 \cdot 10^5 - 3 \cdot 10^9$ cells | Total bacterial cells | Mixed |
| $P$ | $10^7$ cells | - | Total plasmid-carrying cells | Plasmid |

Table S2: Model parameter values used in the independent-donor model and the parameter ranges sampled in the sensitivity analyses.

| Model parameters | Values (SM/LB) | Parameter range | Description | Source |
|---|---|---|---|---|
| $r$ | $1.7$h$^{-1}$ | $0.8 - 2$h$^{-1}$ | Growth rate of MGE-free cells | Determined from growth curve measurements of recipient or donor cells without MGE |
| $r_p$ | See Table S5 | - | Growth rate of plasmid carriers | - with plasmid |
| $r_l$ | See Table S5 | - | Growth rate of lysogens | - with prophage |
| $r_{lp}$ | See Table S5 | - | Growth rate of cells carrying a plasmid and a prophage | - with both |
| $K$ | $10^9$ | - | Carrying capacity | Fitted to CFU counts of all bacterial cells in Figure 1B |
| $\gamma_V$ | $0.01$h$^{-1}$ | - | Virion death rate | [1] |
| $p'$ | $2.5 \cdot 10^{-11}$mL h$^{-1}$ | $10^{-15} - 10^{-8}$h$^{-1}$ | Maximal conjugation rate | Fitted to the increase in $R_P$ cells in Figure 1B |

Table S2: Model parameter values used in the independent-donor model and the parameter ranges sampled in the sensitivity analyses.

| **Model parameters** | **Values (SM/LB)** | **Parameter range** | **Description** | **Source** |
|---|---|---|---|---|
| $p_l'$ | $p' \cdot 0.1$h$^{-1}$ | - | Maximal conjugation rate to lysogenic cells | Adjusted to give a better fit to Figure 1D&S1 |
| $k_p$ | $0.52$ | - | Scale parameter for growth rate dependence of plasmid conjugation | Fitted to Figure 1B |
| $\tau_p$ | $2.2$h | $0.5-3$h | Time for plasmid transfer | Fitted to Figure 1B |
| $\alpha'$ | $10^{-7}$mL h$^{-1}$<br>$10^{-8}$mL h$^{-1}$ | $10^{-12}-10^{-5}$h$^{-1}$ | Maximal adsorption rate of free phage virions to susceptible cells | In agreement with [2, 3] and adjusted to fit Figure 1D& S1 |
| $l'$ | $3 * 10^{-6}$h$^{-1}$ | $10^{-8}-10^{-1}$h$^{-1}$ | Scale for lysogenization rate | Taken from [3] and adjusted to fit Figure 1D& S1 |
| $i$ | $2 \cdot 10^{-5}$h$^{-1}$ | $10^{-8}-10^{-1}$h$^{-1}$ | Induction rate | Taken from [2, 4] and adjusted to fit Figure 1D& S1 |
| $\beta$ | $65$ virions | $30-160$ virions | Burst size | [3] |
| $k_a$ | $0.92$ | - | Scale parameter for growth rate dependence of phage adsorption | Fitted to Figure 1D& S1 |
| $k_2$ | $2$ | - | Rate exponent for growth rate dependence of phage adsorption | Fitted to Figure 1D& S1 |
| $\tau_d$ | $0.3$h | $0.1-0.5$h | Time from infection to establishment of a prophage | [5] |
| $\tau_l$ | $0.8$h | $0.5-1$h | Latent period (time needed for multiplication during the lytic cycle) | Estimated from phage infection experiments |

Table S3: Model variables used in the common-donor model.

| **Model variables** | **Initial values (SM/LB)** | **Description** | **Genetic background** |
|---|---|---|---|
| $D_{LP}$ | $10^6$ cells<br>$10^8$ cells | Plasmid phage donor | Prophage and plasmid |
| $R$ | $10^7$ cells | Recipient cells | MGE-free |
| $R_L$ | 0 | Recipient lysogens | Prophage |
| $R_P$ | 0 | Recipient plasmid cells | Plasmid |
| $R_{LP}$ | 0 | Recipient plasmid lysogens | Prophage and plasmid |
| $V$ | $5 \cdot 10^4$ virions<br>$5 \cdot 10^6$ virions | Free phage virions | - |
| $T$ | $2 \cdot 10^6$ cells<br>$2 \cdot 10^8$ cells | Total bacterial cells | Mixed |
| $P$ | $10^6$ cells<br>$10^8$ cells | Total plasmid-carrying cells | Plasmid |

Table S4: Model parameter values used in the common-donor model. Parameters that were added or changed to improve the fit to empirical data as compared to the independent-donor model are highlighted in red.

| **Model parameters** | **Values (SM/LB)** | **Description** | **Source** |
|---|---|---|---|
| $r$ | $1.7$h$^{-1}$ | Growth rate of MGE-free cells | Determined from growth curve measurements of recipient or donor cells without MGE |
| $r_p$ | See Table S5 | Growth rate of plasmid carriers | - with plasmid |
| $r_l$ | See Table S5 | Growth rate of lysogens | - with prophage |
| $r_{lp}$ | See Table S5 | Growth rate of cells carrying a plasmid and a prophage | - with both |
| $K$ | $10^9$ | Carrying capacity | Fitted to CFU counts of all bacterial cells in Figure 1B |
| $\gamma_V$ | $0.01$h$^{-1}$ | Virion death rate | [1] |
| $p'$ | $2.5 \cdot 10^{-11}$mL h$^{-1}$ | Maximal conjugation rate | Fitted to the increase in $R_P$ cells in Figure 1B |
| $p'_l$ | $p' \cdot 500$h$^{-1}$ | Maximal conjugation rate to lysogenic cells | Adjusted to give a better fit to Figure 3B&C |
| $k_p$ | $0.52$ | Scale parameter for growth rate dependence of plasmid conjugation | Fitted to Figure 1B |
| $\tau_p$ | $2.2$h | Time for plasmid transfer | Fitted to Figure 1B |
| $\alpha'$ | $10^{-7}$mL h$^{-1}$<br>$10^{-8}$mL h$^{-1}$ | Maximal adsorption rate of free phage virions to susceptible cells | In agreement with [2, 3] and adjusted to fit Figure 1D& S1 |
| $\alpha'_p$ | $\alpha' \cdot 10$h$^{-1}$ | Maximal adsorption rate of free phage virions to plasmid carriers | Adjusted to give a better fit to Figure 3B&C |
| $l'$ | $3 * 10^{-6}$h$^{-1}$ | Scale for lysogenization rate | Taken from [3] and adjusted to fit Figure 1D& S1 |
| $i$ | $2 \cdot 10^{-5}$h$^{-1}$ | Induction rate | Taken from [2, 4] and adjusted to fit Figure 1D& S1 |
| $\beta$ | $65$ virions | Burst size | [3] |
| $k_a$ | $0.92$ | Scale parameter for growth rate dependence of phage adsorption | Fitted to Figure 1D& S1 |
| $k_2$ | $2$ | Rate exponent for growth rate dependence of phage adsorption | Fitted to Figure 1D& S1 |
| $\tau_d$ | $0.3$h | Time from infection to establishment of a prophage | [5] |
| $\tau_l$ | $0.8$h | Latent period (time needed for multiplication during the lytic cycle) | Estimated from phage infection experiments |

Table S5: Relative growth rates of recipient or donor cells with various genetic backgrounds, i.e., carrying no, one or both MGEs.

| | **Recipient** $R$ | **Recipient lysogen** $R_L$ | **Recipient plasmid** $R_P$ | **Recipient plasmid lysogen** $R_{LP}/R_{PL}$ | **Donor lysogen** $D_L$ | **Donor plasmid** $D_P$ | **Donor plasmid lysogen** $D_{LP}/D_{PL}$ |
|---|---|---|---|---|---|---|---|
| LB | 1 | 0.63 | 0.73 | 0.76 | 0.89 | 0.92 | 0.89 |
| SM | 1 | 0.93 | 0.73 | 0.79 | 0.94 | 0.90 | 1.1 |
